# Electric fields trigger ceramide-dependent budding of exosome vesicles into multivesicular endosomes and boost the generation of exosomes

**DOI:** 10.1101/2025.06.25.661423

**Authors:** Shiwen Huang, Liangjing Chen, Chuan Sun, Jie Zhang, Tianhui Chen, Yifei Cao, Qin Song, Xianrong Xu, Jianyun Zhang, Xiaohua Tan, Jiandong Wu, Guangdi Chen, Min Zhao, Jun Yang, Yan Zhang

## Abstract

Exosomes hold immense therapeutic and diagnostic potential, yet their clinical translation remains constrained by low yields. Physiological electric fields (EFs), naturally occurring during wound healing and embryo development, have unexplored roles in exosome biogenesis. Here, we demonstrate that physiological-strength EFs (Direct Current, 50-200 mV/mm) dramatically enhance cellular exosome secretion, achieving a nearly 100-fold increase in a strength-dependent manner. Mechanistically, EFs augmented intraluminal vesicle formation within multivesicular bodies. We proposed a novel EF-driven membrane electroassembly and vesiculogenesis hypothesis and tested it using giant plasma membrane vesicles. We reconstructed EF-induced membrane remodeling and observed EFs driving inward budding of nanoscale exosome-like vesicles. Notably, alternating current EFs (50–400 Hz) exhibited significantly diminished efficacy compared to direct current EFs, highlighting membrane electropolarization-dependent modulation. Pharmacological inhibition of the lipid raft and ceramide almost abolishes EF-induced exosome secretion. Inhibition of the PI3K partially attenuated EF-triggered exosome release. Our findings not only unveil EFs as a potent physiological regulator of exosome secretion but also establish a novel, high-yield production strategy leveraging bioelectric cues.

**Significant statement:** We reported that the body’s natural bioelectricity, the faint electric currents in healing wounds and embryo development, can trigger cells to release 100 times more “natural nanoscale vesicle couriers” called exosomes. This pure physical method is far more efficient than current available technologies, paving the way for breakthroughs in cancer early detection, stem cell therapies, and precision medicine. We found that gentle electric fields can “blow” the cell membrane to release exosomes like “soap bubbles”. This explains how bioelectricity controls cell communication during development and wound healing. In short, harnessing the body’s hidden “electric language” unlocks a fast and natural way to mass-produce exosomes.

## Introduction

Exosomes, nanoscale extracellular vesicles (40–150 nm in diameter) secreted by nearly all mammalian cells, are key mediators of intercellular communication. Exosomes have garnered significant attention as cell-free therapeutics, diagnostic tools, and drug-delivery vehicles due to their biomolecular mimicry of parent cells, high biocompatibility, low immunogenicity, and capacity to traverse biological barriers[1, 2]. Despite their clinical promise, the scalable production of exosomes remains a critical bottleneck. Current strategies to enhance secretion — such as 3D culture systems and external stimuli (mechanical stress, hypoxia, chemical agents) — achieve 3- to 20-fold increases in yield, far below the demands of therapeutic or industrial applications [3, 4].

Exosome biogenesis is a tightly regulated process. It begins with plasma membrane invagination to form early endosomes, which mature into late endosomes. Within these compartments, intraluminal vesicles (ILVs) bud inward to generate multivesicular bodies (MVBs), a process mediated by either the endosomal sorting complex required for transport (ESCRT)-dependent or ESCRT-independent pathways (e.g., the nSMase2-ceramide-dependent pathway [5–7]). Mature MVBs either fuse with the plasma membrane to release exosomes or undergo lysosomal degradation [8]. Factors that enhance MVB formation represent a key lever for boosting exosome production.

Endogenous electric fields (EFs), which naturally arise during embryonic development, wound healing, and tumor progression, have emerged as potent regulators of cellular behavior[9–11]. For instance, in skin and corneal wounds, a direct current (DC) EF (25–200 mV/mm) forms at the injury site, directing cell migration and serving as a dominant guidance signal during tissue repair [11–15]. While the mechanisms underlying EF-guided cell migration are partially understood, involving electrophoretic or electroosmosis redistribution of charged membrane components (e.g., integrins, growth factor receptors) within lipid rafts and subsequent activation of signaling pathways such as phosphoinositide 3-kinase(PI3K)/Akt [12, 16, 17]. Previous research has investigated the biological effects of physiological EFs at the cellular and tissue levels; however, the direct interaction between cell membranes and physiological EFs remains largely unexplored. Additionally, the role of physiological EFs in modulating exosome biogenesis is still poorly understood. Here, we systematically investigate the impact of DC EFs at physiological strength on exosome secretion, probing the significance of EF directionality and employing giant plasma membrane vesicles (GPMVs) to dissect EF-membrane interactions. By elucidating how EFs modulate membrane redistribution, ILV budding, and MVB dynamics, this study bridges the gap between bioelectrical signaling and extracellular vesicle biology. Our findings not only unveil physiological EFs as a novel regulator of exosome biogenesis but also establish a scalable, non-invasive strategy for high-yield exosome production, with transformative implications for regenerative medicine and therapeutic delivery.

## Results

### 1. EFs Boost the Production of Exosomes

We utilized a classical galvanotaxis chamber to induce cells to produce exosomes. This system allows a monolayer of cells to adhere to the cell culture dish, be immersed in intensity and direction-controlled EFs, and release exosomes into the culture medium while maintaining cell viability for 8 hours (Figure 1A, Supplementary Figure 1A, B). To examine the effects of EFs on exosome secretion, human skin keratinocytes (HaCaT) were exposed to DC EFs ranging from 0 to 300 mV/mm for 8 hours, followed by nanoparticle tracking analysis (NTA) for quantification. As demonstrated in Figures 1 B, C, the production of exosomes derived from HaCaT cells exhibited a significant increase that depended on the strength of the applied EFs within the physiological range (0-200 mV/mm). Compared to the non-stimulated control, exosome secretion showed a non-significant increase at 50 mV/mm (50 mV/mm vs. Ctrl: 14.61 ± 1.95 vs. 9.86 ± 4.95 exosomes per cell, p > 0.05), but markedly increased at 100 mV/mm (100 mV/mm vs. Ctrl: 410.60 ± 27.75 vs. 9.86±4.95, exosomes per cell, p < 0.01), peaking at 200 mV/mm with a more than 100-fold increase in secreted exosomes compared to the control (200 mV/mm vs. Ctrl:1361.00 ± 203.10 vs. 9.86 ± 4.95 exosomes per cell, p < 0.01)(Figure 1 B, C). The vesicle size distribution remained comparable to that of the control group (Figure 1D, Supplementary Figure 1C). Additionally, Western blot analysis indicated higher levels of Alix, CD63, CD9, and CD81 in the exosomes produced by cells stimulated at 200 mV/mm compared to the control group (Figure 1E). A slight decline in exosome production was observed at 300 mV/mm, suggesting that electric fields exceeding the physiological range do not further enhance exosome secretion (Figure 1B, C). Post-stimulation tracking revealed a rapid decrease in exosome production after the withdrawal of the EFs (Figure 1F). Notably, physiological EFs demonstrated superior efficacy compared to other conventional exosome-inducing stimuli, such as starvation and heat treatment (Figure 1G). Validation in human embryonic stem cell-derived mesenchymal stem cells (MSCs) and HT1080 fibrosarcoma cells confirmed the overarching capacity of EFs to enhance exosome secretion. (Figure 1I-L, Supplementary Figure 1G-J). In contrast, physiological EFs did not significantly promote the secretion of extracellular microvesicles (MVs) (Figure 1H, Supplementary Figure1D-F). Collectively, these data demonstrate that physiological-strength EFs substantially enhance exosome secretion across various cell types.

**Figure 1.**
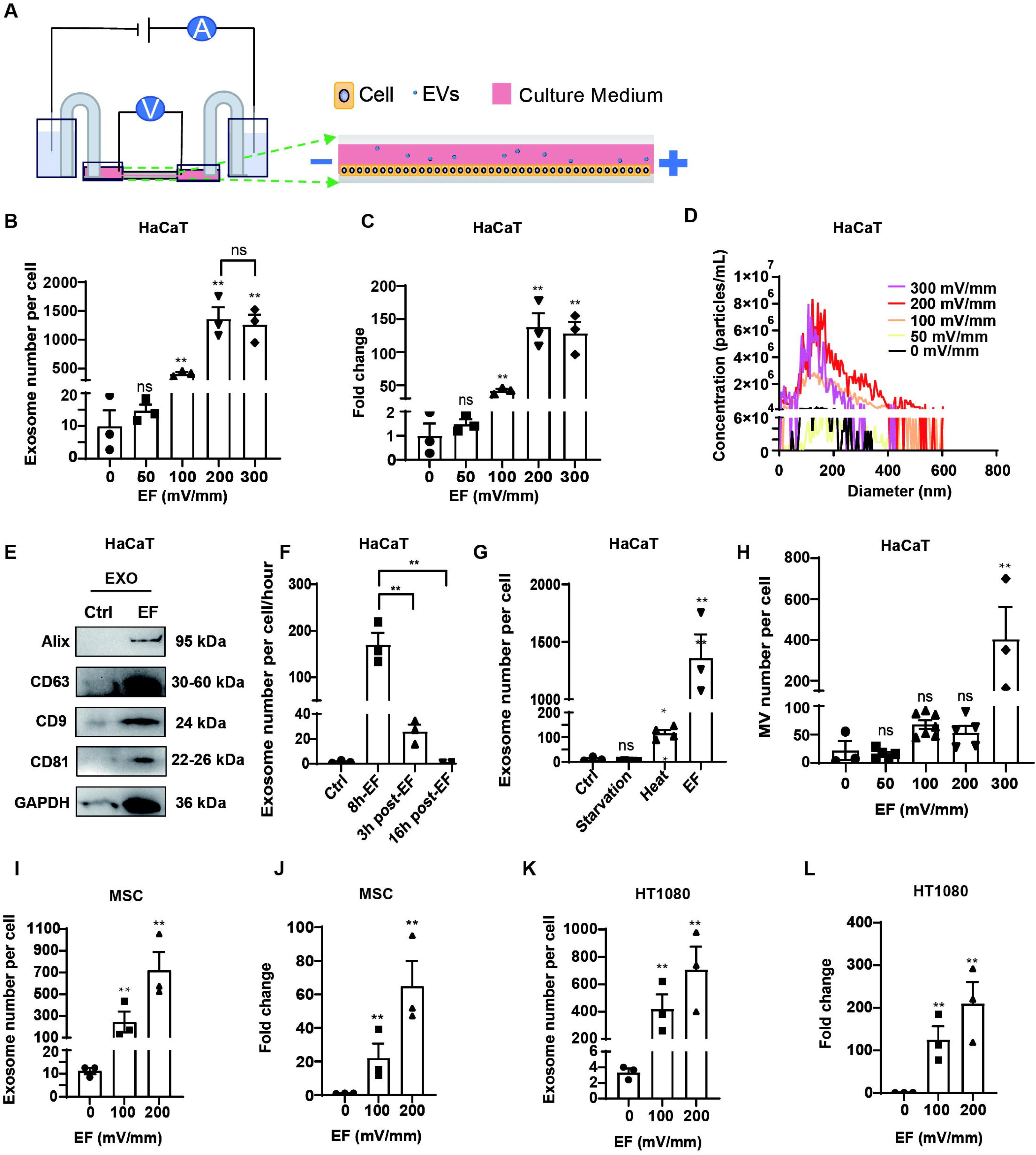
EFs boost the generation of exosomes. (A) A schematic representation of the electric stimulation device. A monolayer of cells was seeded in the central area of the galvanotaxis chip, releasing extracellular vesicles (EVs) into the cell culture medium. (B–D) NTA results showing: (B) the number of exosomes per cell; (C) the fold change in the number of exosomes; and (D) the diameter distribution of exosomes produced by HaCaT cells under different EF strengths over 8 hours. (E) Western blot results demonstrating differing expression levels of Alix, CD63, CD9, CD81, and GAPDH in exosomes from the control (no EF) group and the EF (200 mV/mm) group, measured at 8 hours. (F) NTA results showing the exosome count per cell per hour in the control group, the EF (200 mV/mm, 8 hours) group, and at 3 and 16 hours post-EF stimulation groups. (G) Comparison of EV release using the EF method versus traditional approaches for stress-induced exosome release, including starvation and heat treatment. For starvation, HaCaT cells were cultured in DMEM lacking glucose and FBS for 8 hours. For heat stress, HaCaT cells were incubated at 42 ℃ for 2 hours before being transferred to 37 °C under normal cell culture conditions for 6 hours. (H) NTA results showing the number of extracellular microvesicles (MVs) per cell produced by HaCaT cells under physiological EFs (50-200 mV/mm, 8 h) didn’t increase significantly, but increased at 300 mV/mm. (I–L) The exosome counts per cell and the fold change produced by MSC and HT1080 cells under different EF strengths over 8 hours. All data were derived from three independent experiments and are presented as means ± standard error of the mean (SEM). ns indicates not significant, *p < 0.05, **p < 0.01.

### 2. EF-induced MSC-derived exosomes retain native capacities for genomic safeguarding and cellular function promotion

To evaluate the biological efficacy of exosomes derived from EF-induced exosome (Exo^EF^) derived from MSCs, we compared their DNA-protective capacity against cisplatin (Pt)-induced genotoxicity with conventionally produced exosomes (Exo^Ctrl^). Using comet assays, HaCaT cells pretreated with Exo^EF^ or Exo^Ctrl^ (harvested from equal quantities of 8-hour MSC secretions) showed distinct responses to Pt (10 µM, 24 h). As demonstrated in Figure 2A-D, Pt treatment resulted in significant DNA damage, as indicated by the presence of elongated comet tails, characterized by increased percentages of tail DNA, tail moment, and Olive tail moment in the Pt-only group. Exo^EF^ pretreatment significantly attenuated these metrics, reducing comet tail parameters to levels indistinguishable from Pt-untreated controls. Notably, Exo^EF^ outperformed Exo^Ctrl^, suggesting enhanced intrinsic DNA-protective activity.

**Figure 2.**
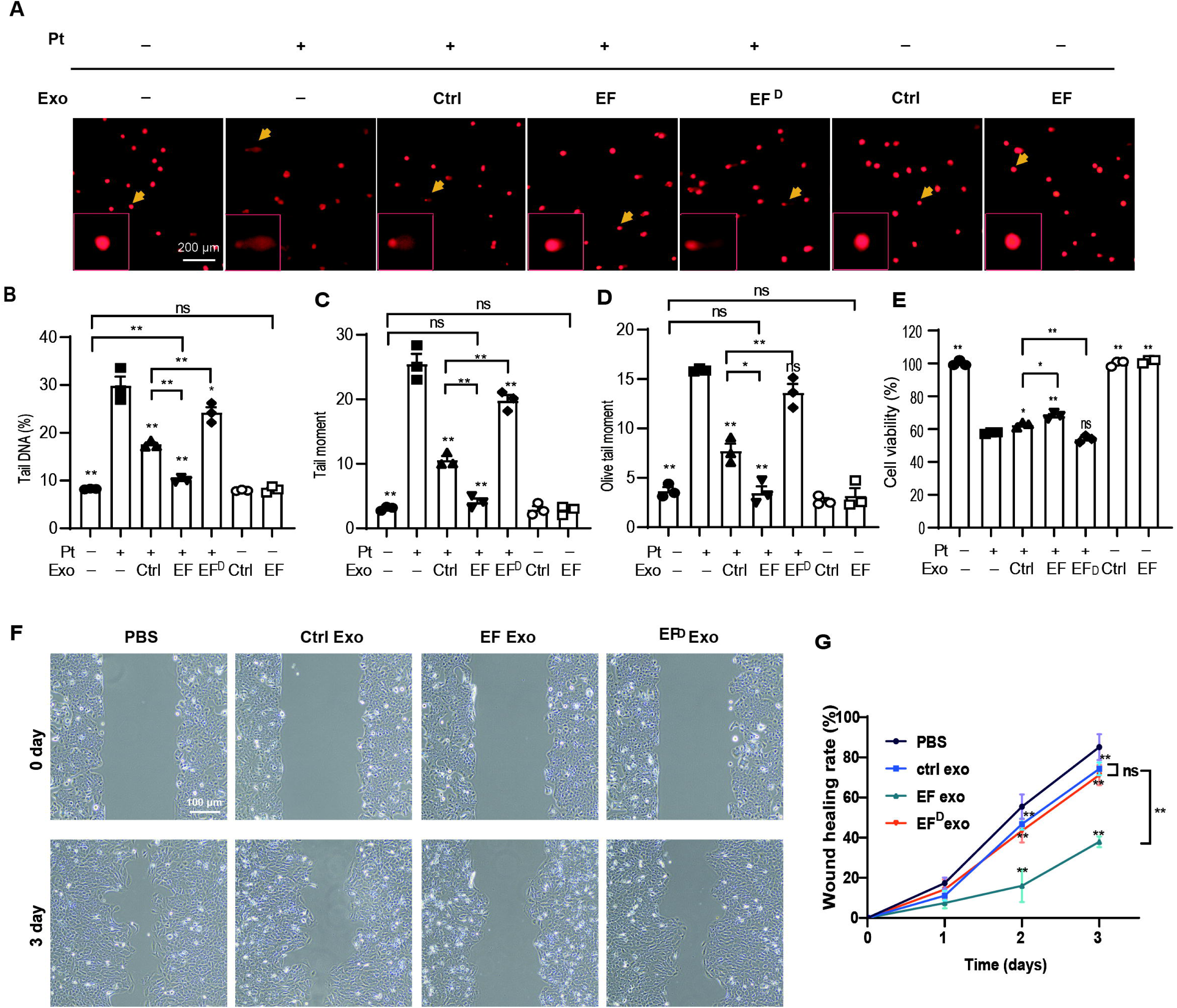
Function analysis of EF-induced MSC exosomes. (A) Representative comet assay images showing DNA fragmentation in HaCaT cells after different treatments. Cells were treated with PBS, MSC-derived exosomes from the control group, EF group, and EF^D^ group for 6 hours, followed by the addition of 10 µMol/L Pt for another 24 hours. EF^D^ group indicates cells treated with the exosomes of the EF group (200 mV/mm, 8 hours) (×80) diluted to the same concentration as the non-stimulated control group (5×10^5^ particles/mL). (B-D) Tail DNA (%) (B), tail moment (C), and olive tail moment (D) were analyzed. More than 100 cells per group were measured in each experiment. Compared to the Pt-only treatment group. (E) Cell viability of each group was also measured by trypan blue cell counting assay. (F-G) Wound healing assay of HaCaT cells with different treatments as previously described. Data are presented as mean ± SEM from three independent experiments. For A-E, unless otherwise stated, compared with the Pt-only group. For E, G, unless otherwise stated, compared with the control group. ns indicates not significant, *p < 0.05, **p < 0.01.

However, when Exo^EF^ was diluted to match the concentration of Exo^Ctrl^ (5 × 10^5^ particles/mL)(Exo^EFD^), the outcomes diverged. (Figure 2A-D). Under iso-concentration conditions, Exo^EFD^-pretreated cells exhibited longer comet tails than Exo^Ctrl^-pretreated cells, implying reduced per-particle efficacy. This paradox highlights that the superior performance of Exo^EF^ at matched cells arises from increased exosome yield under electrical stimulation, rather than enhanced bioactivity per vesicle. Similar results were observed in cell viability assay (Figure 2E) and a wound healing assay using HaCaT cells (Figure 2F, G). Consistent with previous reports [18], exosomes derived from MSCs impaired wound healing in the scratch assay, and Exo^EF^ has a more pronounced inhibitory effect (Figure 2F, G). Collectively, physiological electrical preconditioning amplifies MSC exosome secretion without compromising intrinsic DNA protection and cellular function regulation capacity.

### 3. EFs promote the generation of ILVs in MVBs

The formation of MVBs represents a critical step in exosome biogenesis. To investigate whether physiological EFs modulate exosome secretion by influencing MVBs generation in HaCaT cells, we first analyzed MVB ultrastructure and abundance using transmission electron microscopy (TEM). Morphological examination revealed that both EF-treated and untreated groups exhibited MVBs with the characteristic “vesicles-in-a-sac” architecture, containing numerous intraluminal vesicles (ILVs) in the cytoplasm (Figure 3A). Quantitative analysis demonstrated no significant difference in MVB density per cellular cross-sectional area between groups (Figure 3B). Strikingly, EF stimulation markedly increased ILV cargo load within individual MVBs compared to controls (Figure 3C). PKH26 membrane labeling combined with fluorescence microscopy analysis showed comparable surface fluorescence intensity between groups (Figure 3D), further suggesting unchanged bulk membrane trafficking activity. These findings collectively indicate that physiological EFs promote exosome secretion by specifically enhancing ILV packaging efficiency within MVBs. This EF-triggered amplification of ILV synthesis suggestes a mechanistic basis for the observed exosome secretion potentiation.

**Figure 3.**
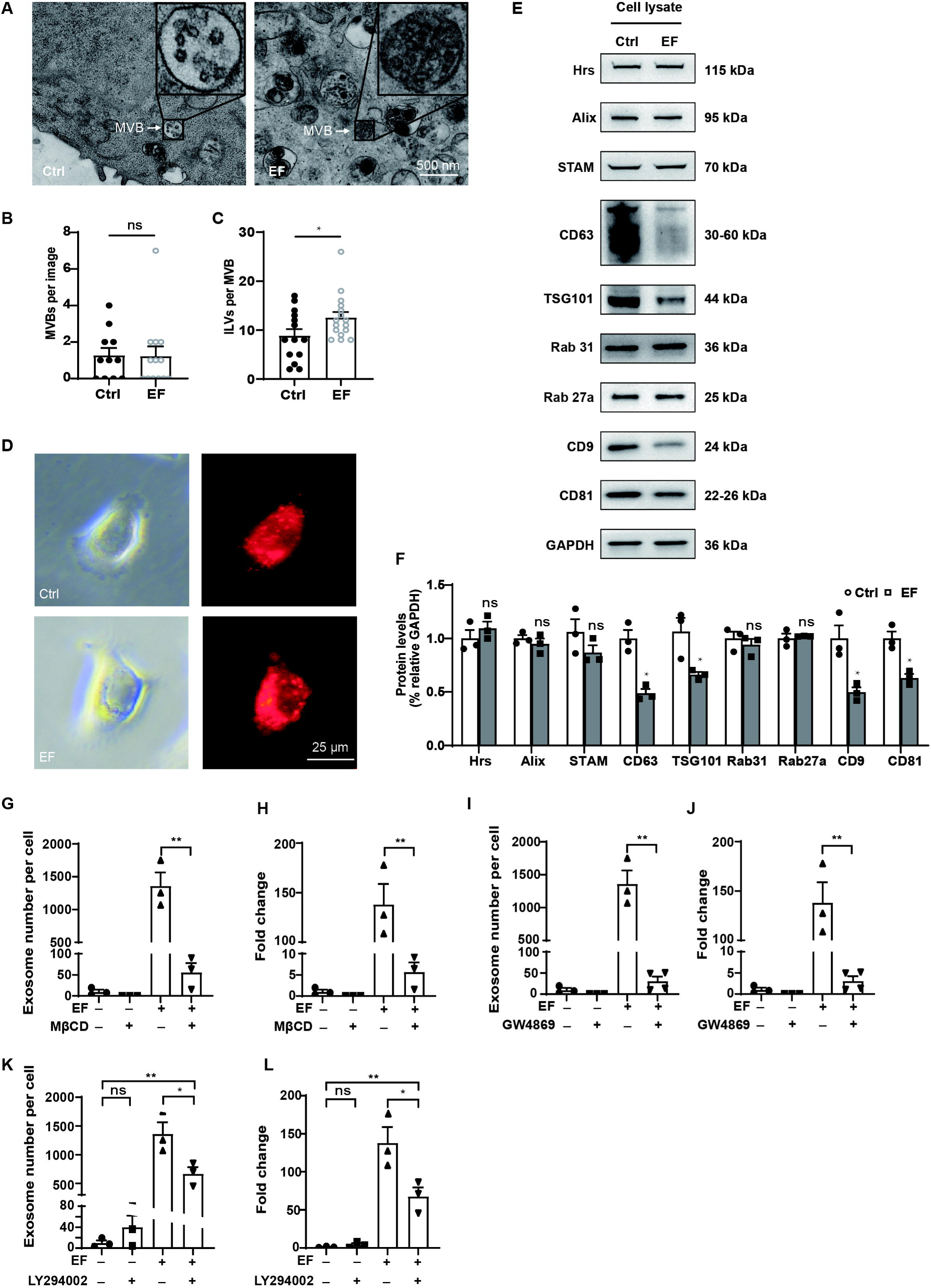
EFs stimulate exosome biogenesis mainly by ESCRT-independent mechanisms. (A) Representative TEM images demonstrate EF-treated HaCaT cells (200 mV/mm, 8 hours) exhibit increased ILVs within MVBs compared to untreated control cells. (B, C) Quantitative analysis of TEM images reveal no significant change in MVB counts per cell section (B) and significant elevation of ILVs per MVB profile (C). Data derived from >10 independent TEM fields per group. (D) Fluorescence images show no significant change in intracellular vesicle formation in HaCaT cells between control and EF groups, measured as spots of red fluorescence from PKH26 dye. (E, F) Western blot identifies the EF-induced downregulation of CD63, TSG101, CD9, and CD81, with concomitant no significant change in expression of ESCRT-related proteins. (G-L) Disrupting lipid rafts with MβCD (4 mM) (G, H) or inhibiting neutral sphingomyelinase 2 activity with GW4869 (10 μM) (I, J) significantly reduced EF-induced exosome release in HaCaT cells. Inhibiting PI3K with LY294002 (35 μM) partly inhibited the release of exosomes induced by EFs. Data are presented as mean ± SEM from three independent experiments. Unless otherwise stated, compared with the control group. ns indicates not significant, *p < 0.05, **p < 0.01.

### 4. EF-triggered exosome secretion requires functional lipid raft and ceramide

To further dissect the mechanistic basis of EF-induced exosome biogenesis, we analyzed protein expression profiles associated with distinct exosome generation pathways using Western blot (Figure 3E). Quantitative assessment of ESCRT-complex components (Hrs, STAM) and Rab31 (inhibitor of MVB-lysosome fusion), Rab27a (promoter of MVB-plasma membrane docking), and Alix (scaffold protein for vesicle budding)[8] showed comparable expression levels between groups. In contrast, exosome protein markers, including TSG101, CD9, CD63, and CD81, exhibited reduced expression in EF-treated cells (Figure 3F). These findings collectively demonstrate that physiological EFs promote exosome generation primarily through an ESCRT-independent pathway. The selective depletion of TSG101 and tetraspanins may reflect enhanced exosomal expulsion rather than impaired production, consistent with our prior observation of EF-triggered exosome secretion amplification.

To test the hypothesis that physiological EFs promote exosome secretion via lipid raft-dependent pathways, we disrupted cholesterol-rich membrane microdomains using methyl-β-cyclodextrin (MβCD, 4 mM, 1 hour) and inhibited generation of ceramide by inhibiting neutral sphingomyelinase 2 activity with GW4869 (10 μM), both critical for lipid raft integrity and ESCRT-independent exosome biogenesis. Strikingly, EF-induced exosome release in HaCaT cells was almost abolished by either inhibitor (Figure 3G-J). Quantitative NTA demonstrated that MβCD pretreatment suppressed exosome secretion in the 200 mV/mm electric field group, reaching only 4.12 ± 0.11% of the EF-only group. Similarly, GW4869 decreased exosome production to 2.26 ± 0.05% of the EF-only levels (Figure 3G-J). These findings demonstrate that the nSMase2-ceramide-dependent pathway is an essential prerequisite for EF-driven exosome release.

Physiological EFs can activate the cellular phosphoinositide 3-kinase (PI3K)/Akt pathway, while the inhibition of this pathway significantly impairs directional cell migration induced by EFs [12]. To investigate whether PI3K/Akt also plays a role in exosome biogenesis triggered by EFs, we inhibited PI3K using LY294002. Our results from NTA indicated that the stimulatory effect of EFs on exosome secretion was partially diminished (Figure 3K, L). This finding suggests that the signaling pathways involved in EF-induced exosome secretion and EF-guided cell migration are at least partially overlapping, though additional mechanisms may also govern the regulation of exosome secretion triggered by EFs.

### 5. Polarization persistence governs secretion efficiency

Drawing upon the intrinsic capacity of membrane molecules (e.g., cone-shaped ceramide) aggregation and self-assembly to generate vesicles[8], as well as the observation that physiological EFs can polarize charged molecules on membranes[8, 16, 19] and increase the number of ILVs in MVBs within EF-treated groups (Figure 3A), we propose a novel hypothesis: the EF-Driven Membrane Electroassembly and Vesiculogenesis (EF-MEV) hypothesis. This hypothesis posits that EFs may enhance the aggregation and self-organization of charged molecules on membranes, thereby facilitating the inward formation of vesicles and ultimately promoting the production of exosomes (Figure 4A).

**Figure 4.**
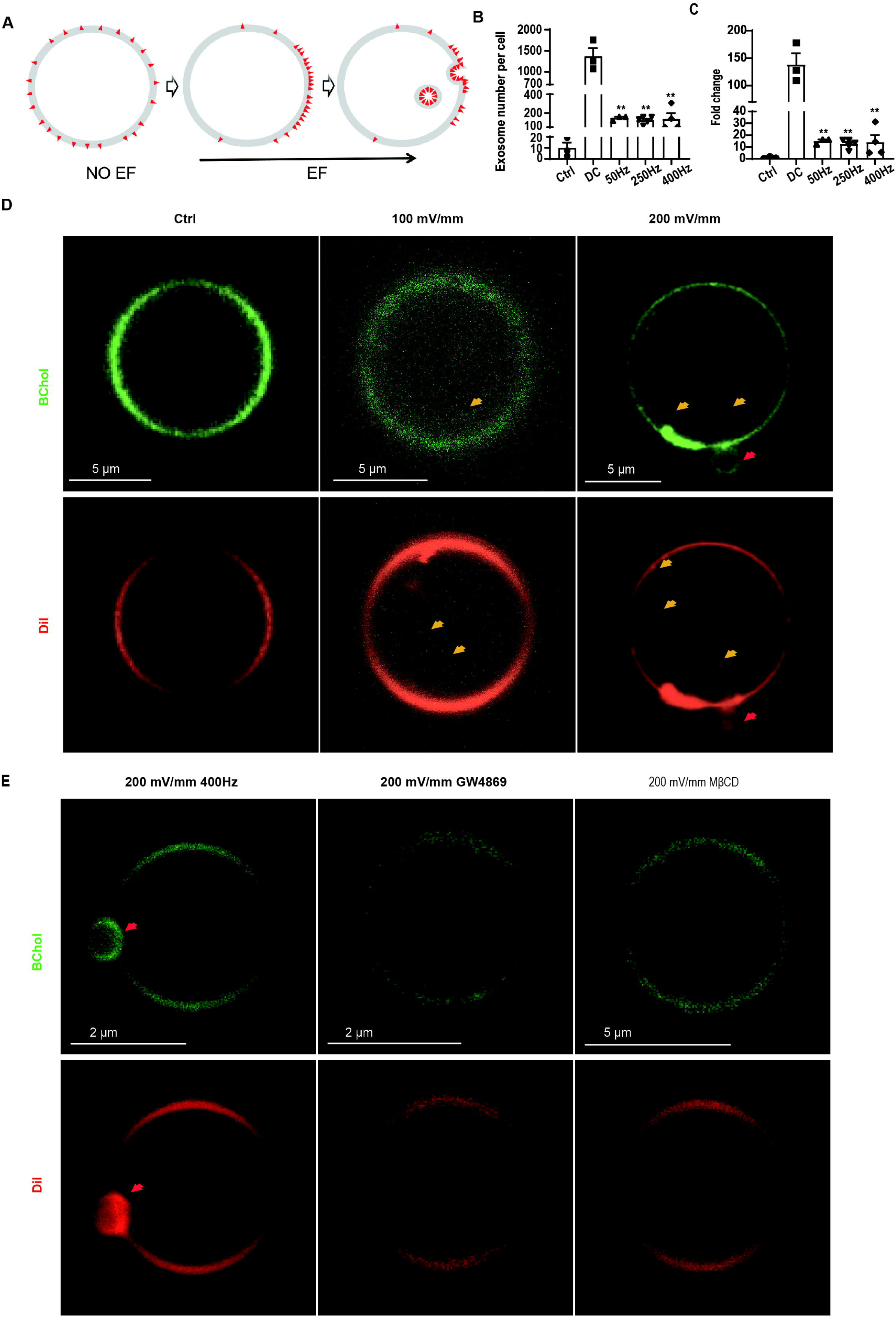
EFs trigger the budding of exosome-like vesicles into GPMVs. (A) Hypothetical schematic of EF-induced membrane vesiculation (EF-MEV) hypothesis. DC EFs promote lateral migration and aggregation of charged conical lipids (red triangles) within lipid bilayers by electrophoresis and/or electroosmosis. This molecular rearrangement generates intrinsic membrane curvature, driving the formation of nano-vesicles that bud inward to form exosome precursors. Note: Vesicle sizes are not drawn to scale. (B, C) Quantification of exosome production and the fold change in the number of exosomes under AC EF stimulation. NTA revealed a decrease in exosome yield per HaCaT cell following AC EF exposure (200 mV/mm, 50-400 Hz, 8 hours) compared to the DC EF group (200 mV/mm, 8 hours). (D-E) Visualization of EF-induced vesiculation in GPMVs. Confocal microscopy of DiI(red)/BChol(green)-labeled GPMVs exposed to DC/AC EFs (0∼200 mV/mm, 0∼400Hz, 2 hours) revealed spontaneous generation of nano-vesicles (orange arrows) in DC EF-treated GPMVs. The nanoscale vesicles within GPMVs exhibit rapid Brownian motion under microscopy observation. This results in motion-induced blurring when captured with prolonged exposure times (typically >50 ms). Furthermore, since GPMVs remain freely suspended in an aqueous medium without complete immobilization during imaging, sequentially acquired fluorescence micrographs cannot achieve perfect spatial superposition. No detectable nano-vesicles were observed in GPMVs derived from cells pretreated with GW4869 (10 μM, 10 hours) and MβCD (4 mM, 1 hour). Sporadic micron-scale vesicles budding outward from GPMVs (red arrows) were also observed in the DC and AC EF groups. Data were presented as mean ± SEM of three independent experiments. Unless otherwise stated, compared with the control group. *p < 0.05, **p < 0.01.

To validate this hypothesis, we first investigated whether the EFs promote cell secretion in a manner dependent on molecular polarization on the membrane. High-frequency alternating currents (AC) can inhibit the polarization of charged molecules in EFs [16]. We applied an 8-hour alternating EF of 200 mV/mm at various frequencies (50-400 Hz) to HaCaT cells to dynamically disrupt the establishment of membrane polarization steady state. As shown in Figure 4B, C, the amount of exosome secretion in the AC EF group was significantly lower than that in the 200 mV/mm DC EF group, decreasing to 10.87 ± 0.07% to 19.78 ± 0.68% of the DC group. These findings suggest that the maintenance of a steady state of membrane polarization is a necessary condition for physiological EFs to promote exosome secretion.

### 6. EFs trigger the budding of exosome-like vesicles into GPMVs

To directly validate our EF-MEV hypothesis, we analyzed the effects of physiological EFs on membrane dynamics using GPMVs. These cell-derived micron-scale vesicles retain native membrane composition and protein content while eliminating confounding cellular processes (e.g., protein synthesis, active transport, cytoskeletal interactions, and signaling cascades) [20, 21]. This makes them especially well-suited for exploring membrane-specific phenomena related to the interactions between EFs and cellular membranes.

We exposed DiD-C18 and BODIPY-Cholesterol-stained GPMVs to 2-hour stimulation with DC or AC EFs. Confocal imaging revealed a striking disparity between treatment groups: about 30% of GPMVs exposed to 200 mV/mm DC fields exhibited abundant inward budding of exosome-like nanoscale vesicles, accompanied by stained membrane aggregation on one side of the GPMV, whereas unstimulated controls and AC field-treated groups showed no significant nanoscale vesiculation (Figure 4D). The above observations in GPMVs in response to EF aligned with reported characteristics of exosome biogenesis in EFs (Figure 3A, C).

To identify critical membrane components mediating this response, we disrupted lipid rafts using MβCD (4 mM, 1-hour pretreatment). MβCD-treated cells yielded smaller GPMVs that failed to generate nano-vesicles under DC field stimulation (Figure 4E). Subsequent pharmacological inhibition of ceramide synthesis using GW4869 (10 μM, 10-hour pretreatment), similarly abolished field-induced vesiculation, with ceramide-depleted GPMVs showing complete loss of inward budding (Figure 4E). The above results further support our EF-MEV hypothesis and suggest EF-MEV is ceramide-dependent.

Intriguingly, we also observed sporadic micron-scale vesicles budding inward (Supplementary Figure 2A) and outward (Figure 4D, E) from GPMV membranes in control, DC and AC EF groups. Quantitative analysis revealed a significant increase in these membrane protrusions in DC EFs (200 mV/mm, 2 hours) compared to controls. However, three critical findings suggested non-exosomal origins: 1) GW4869 (10 μM) failed to reduce vesicle counts; 2) switching to AC fields (400Hz) maintained elevated vesicle density; 3) the dimensions of these structures significantly surpass those of exosomes (Supplementary Figure 2A, B). Based on these pharmacological and biophysical evidence, we concluded the phenomenon represents a distinct membrane remodeling process unrelated to classic exosome biogenesis pathways.

## Discussion

Our findings establish that physiological DC EFs enhance exosome secretion through a lipid raft-dependent mechanism involving membrane electropolarization and ceramide-mediated curvature generation. This work bridges two previously distinct domains—bioelectric signaling and membrane biophysics — to propose a unified framework for EF-driven exosome biogenesis.

### 1. EFs enhance exosome secretion in a quantity-over-quality paradigm

Our findings demonstrate that physiologically relevant EFs significantly enhance exosome secretion in a strength-dependent manner (Figure 1B, C; I-L), with secretion levels rapidly declining upon EF withdrawal (Figure 1F). This transient effect contrasts sharply with existing electrical stimulation strategies, such as high-voltage electroporation [22] and iontophoresis [23], which rely on supraphysiological intensities to achieve exosome release through thermal and mechanical disruption of membranes. While these high-intensity methods yield high exosome quantities, their cytotoxic effects and non-physiological mechanisms limit clinical applicability [22]. In contrast, our 200 mV/mm DC EFs, mimicking endogenous wound-edge EFs, operate within physiological parameters and preserve cell viability during prolonged stimulation (Figure S1A, B). Compared to biochemical inducers (e.g., hypoxia or starvation), EFs boosted secretion by nearly 100-fold (vs. reported 3–20× increases for conventional stimuli) (Figure 1G), underscoring their scalability potential. Together, this positions physiological EFs as a biocompatible alternative for sustained exosome production.

The preserved DNA-protective and cellular function regulation capacity of EF-induced exosomes—despite a nearly 100-fold increased secretion—further supports a quantity-over-quality paradigm, where EFs boost vesicle quantity without altering intrinsic functionality (Figure 2A-G). The reduced per-particle efficacy of Exo^EF^ at iso-concentration (Figure 2A-G) cautions against assuming enhanced bioactivity—EFs act as secretion amplifiers, not molecular potentiators. This duality—enhanced secretion but variable potency, highlights the need for comprehensive cargo profiling (e.g., proteomic/RNA sequencing) to optimize therapeutic exosome harvest. Similar results have been reported regarding mesenchymal stem cell-derived small extracellular vesicles induced by short-term small EFs (100 mV/mm for 1 hour per day), which enhance neuroprotective effects in spinal cord injury. However, due to the brief duration of stimulation, the study did not detect an increase in exosome production [24].

### 2. EFs drive membrane electroassembly and vesiculogenesis

The EF-driven exosome hypersecretion operates via an ESCRT-independent pathway, as evidenced by unaltered ESCRT complex proteins and Rab GTPases regulating vesicle trafficking (Figure 3E, F). Instead, EF efficacy critically depended on lipid raft integrity and ceramide synthesis, with MβCD and GW4869 abolishing the >95% of EF-induced secretion (Fig. 3G-J). These findings align with ceramide’s established role in ESCRT-independent ILV formation within MVBs, where its conical geometry induces negative membrane curvature to bud inward vesicles [7].

We propose a novel EF-DMEV model (Figure 4A) in which physiological EFs polarize charged lipid raft components, such as cone-shaped structures of ceramide, through electrophoresis or electroosmosis, promoting their aggregation at the membrane, facilitating the spontaneous induction of negative curvature in the membranes of endosomes and promoting the formation of internal vesicles within MVBs. Supporting this, AC EFs (which prevent stable charge polarization) reduced secretion by >80% (Figure 4B, C), consistent with prior observations of DC-specific bioeffects of electrotaxis [16]. Crucially, EF-induced ILV enrichment within MVBs (Fig. 3A, B) without increasing MVB numbers suggests EFs amplify cargo packaging efficiency rather than initiating MVB biogenesis de novo. The GPMV experiments provided direct evidence of EF-triggered membrane self-assembly (Figure 4D). EFs induced polarized aggregation of membrane and inward nanovesiculation in GPMVs, replicating ILV formation in cells, while raft/ceramide disruption abolished this effect (Figure 4E). This parallels material science paradigms where external EFs guide molecular alignment and self-organization of nanoparticles [25]. Cell membranes, as naturally evolved “smart materials”, appear exquisitely optimized for field-responsive remodeling.

While our GPMV experiments effectively decouple membrane biophysics from cellular complexity, they cannot completely replicate the interaction between EFs and late endosomes. The limitation arises from the differences in membrane components and pH environment between the GPMVs and late endosomes [26]. Additionally, the membrane composition of individual GPMVs may vary throughout generation process. Those variations may account for the observation that only a fraction of GPMVs respond to EFs and produce exosome-like vesicles.

At which stage do EFs interact with membranes to generate exosome-like vesicles during process of exosome generation? Theoretically, the resistance of the cell membrane is exceptionally high, akin to a “Faraday cage,” which tends to shield EFs from penetrating the interior of the cell. However, there are reports indicating that ion flow can occur within the cell body in the presence of EFs due to the opening of ion channels [27]. Therefore, the interaction between EFs and the membrane could take place at the cell membrane, leading to aggregation. Nanovesiculation, on the other hand, may occur at the stage of late endosomes. Alternatively, EFs might directly interact with the membranes of late endosomes, facilitating the generation of nanovesicles.

### 3. Future directions

While PI3K inhibition partially attenuated EF-induced secretion (Figure 3K, L), its ∼50% efficacy implies that parallel activated pathways previously reported involved in directional migration in EFs (e.g., Ca²^+^ influx [28] or ROS signaling [29]) might also contribute to exosome regulation. PI3K [30], Ca²+ influx [22] have already been demonstrated to be involved in exosome regulation. The interplay between EF-directed cell migration and exosome secretion also warrants investigation.

Our TEM and GPMV experiments suggest that EFs could initiate ceramide-dependent budding of exosomal vesicles into MVBs (Figure 3A, Figure 4D). However, while the activated signaling pathways in response to EFs may also contribute to the increase in exosome secretion, quantifying the precise contribution of this phenomenon to exosome release remains challenging. Building upon elegant giant unilamellar vesicles (GUVs) studies demonstrating pH-dependent lysobisphosphatidic acid microdomain assembly [26], ESCRT-III and Vps4 coupling [31, 32] during the formation of MVBs, our future work will employ GUVs with precisely controlled lipid compositions and transmembrane pH gradients, contingent on our comprehensive understanding of the lipids, forces, and microenvironment necessary for this response. The biological implications of the increase in microscale vesicle number observed in the EF-stimulation group require further investigation (Supplementary Figure 3A).

Our bioelectric field chamber successfully maintained stable, sustainable EFs and cell viability for up to 8 hours (Supplementary Figure 1A, B); however, cellular responses to EF stimulation beyond this duration have not yet been investigated. Consistent with previous reports [33], EF stimulation depletes membrane components, including CD9, et al. (Figure 3E). Further research is necessary to determine whether prolonged exposure to EFs could establish a new equilibrium between membrane component synthesis and exosome secretion rates.

The significant effects of EFs on exosome secretion suggest that emerging EFs might play a crucial role in regulating wound healing, embryonic development, and cancer metastasis. Although the guidance ability of EFs on cell migration has been demonstrated in various in vivo models [9, 34, 35], the role of EFs in regulated exosome secretion requires independent investigation in vivo in future studies.

In conclusion, physiological EFs unlock a non-invasive, biomimetic strategy for scalable exosome production while offering unprecedented insights into membrane nanoengineering. By bridging the fields of cell biology and materials science, this work illuminates the untapped potential of bioelectricity in regenerative medicine and nanotechnology.

## Materials and Methods

### Cell cultures

HaCaT, human embryonic stem cell-derived MSCs, and HT1080 cell lines from Yuansheng Biotechnology, Inc. (Hangzhou, China) were cultured in Dulbecco’s modified Eagle medium (DMEM; cat. no. C3113-0500, VivaCell, China) supplemented with 10% fetal bovine serum (FBS; cat. no. NB500-S, NobleBio, AUS) and 1% penicillin–streptomycin (PS; cat. no.15140122, Gibco, USA) at 37 ℃ in humidified conditions equilibrated with 5% CO2. For MSCs, only four to seven generations of cells were selected for the experiment.

### Electrical stimulation

As previously described [36], the galvanotaxis chamber was a modified parallel-plate flow chamber with agarose salt bridges connecting the medium’s entrance and outflow. Constant DC EF was applied at a field strength of 0-300 mV/mm with a DC power supply (Beijing Liuyi DYY-6C, China), and Square-wave AC signals were generated using a programmable AC signal generator (YP4050, YongPeng Instruments; 500 VA, 0–300 V output range, China) with frequency modulation spanning 0–400 Hz.

### Collection of extracellular vesicles (EVs)

To achieve about 80% confluency after an overnight incubation period, HaCaT, MSCs, or HT1080 cells were seeded into the galvanotaxis chamber of 100-mm plates at a density of 4, 2 or 4 × 10^5^ cells/dish, respectively. Following the removal of the medium, 1 phosphate-buffered saline (PBS; cat. no. 10010023, Gibco, USA) was used to wash the cells. Cell culture medium was replaced with 3 mL DMEM supplemented with 2% 1 M HEPES (cat. no. 15630080, Gibco, USA) and 1 % PS. The conditioned medium was collected after electrical stimulation, and EVs were purified from medium via differential centrifugation. Briefly, the medium was subjected to sequential centrifugation at 300 × g for 10 min to remove cells, 2000 × g for 10 min to clear subcellular debris, and 10,000 × g for 30 min to separate microvesicles at 4 ℃, followed by filtration with 0.22-µm syringe filters (cat. no. BS-PES-22, Biosharp, AUS). Then, the samples were added to a 100 KD ultrafiltration tube (cat. no. UFC910096, Millipore, USA) and centrifuged several times at 4,000 × g for 15 min at 4 ℃ to concentrate the exosomes. Next, half the volume of the total exosome isolation reagent (cat.no.4478359, Invitrogen, USA) was added to the concentration solution and left at 4 ℃ overnight. The following day, the exosomes were precipitated by ultracentrifugation (10,000 × g, 60 min, 4 ℃), and the supernatant was aspirated. The exosomes were then resuspended in PBS and used in the following experiments.

### Measurement of EV particle size distribution and particle number

The number and size distribution of particles in the collected EV suspension were measured with a nanoparticle multianalyzer (Zetaview-PMX120-Z, Particle Metrix, GER). The EVs were diluted to 10^7^ particles/mL for NTA.

### Transmission electron microscopy

Cells for TEM analysis were collected with a cell scraper after 8 hours of electrical stimulation, and after centrifugation to aspirate the supernatant, the cells were resuspended in 4% paraformaldehyde and layered on a 200-mesh copper grid, which was subsequently stained with 1% phosphotungstic acid. Grids were air-dried, and intracellular MVBs were observed using an HT-7700 transmission electron microscope.

### Cell staining

Cells in the galvanotaxis chamber were stained with PKH26 using PKH26 Red Fluorescent Cell Linker Kits for General Cell Membrane Labelling (cat. no. PKH26GL-1KT, Sigma-Aldrich, GER). 4 µL of PKH26 dye was diluted in 2 mL of diluent C, and cells were incubated for 5 min. The staining was then terminated by adding 2 mL of FBS and incubating for 1 min. The dye was aspirated, and the cells were rinsed 5 times with 1 mL DMEM. 3 mL of serum-free medium was added, and cells were given 3 hours of DC stimulation. Then, fluorescence was observed and recorded on a fluorescence microscope (ECLIPSE Ts2R, Nikon, Japan).

### Cell viability assay

HaCaT cells were seeded in a 96-well plate with 10,000 cells per well and treated with Exo^Ctrl^ or Exo^EF^ for 6 hours. Then, the cells were treated with serum-free DMEM with DMSO or 14 µM cisplatin (Pt; cat. no. GC11908, GlpBio, USA) for 24 hours. Cell viability was assessed by CCK-8 kit (cat. no. GK10001, GlpBio, USA) according to the manufacturer’s instructions. Additionally, the cells were trypsinized after 8 hours of DC stimulation at different intensities, and 0.4% trypan blue (cat. no. GF00376, GlpBio, USA) staining was used to count the cells.

### Western blotting

After being separated on 10% SDS-PAGE gels (cat. no. ET15010Gel, ACE, China), protein samples were transferred onto PVDF membranes (cat. no. 03010040001, Sigma-Aldrich, USA). The membranes were blocked with 5% nonfat milk (cat. no. A600669-0250, Sangon, China) in 1× Tris-buffered saline solution (TBS; cat. no. ST661-500mL, Beyotime, China) and incubated with primary antibodies in TBS with 0.05% Tween 20 (cat. no. ST825-100mL, Beyotime, China) overnight at 4 ℃. Following the washing step, the blots were reacted with secondary antibodies for 1 hour and developed using the enhanced chemiluminescence (ECL; cat. no. P0018AS, Beyotime, China) detection system. Alix mouse monoclonal antibody (cat. no. sc-53540), CD81 mouse monoclonal antibody (cat. no. sc-23962), CD9 mouse monoclonal antibody (cat. no. sc-18869), CD63 mouse monoclonal antibody (cat. no. sc-5275), Rab27A mouse monoclonal antibody (cat.no.sc-74586), Rab31 mouse monoclonal antibody (cat. no. sc-517069), STAM mouse monoclonal antibody (cat. no. sc-133093), Hrs mouse monoclonal antibody (cat. no. sc-271455), caveolin-1 mouse monoclonal antibody (cat. no. sc-53564), TSG101 mouse monoclonal antibody (cat.no.sc-7964), and GAPDH mouse monoclonal antibody (cat. no. sc-47724) were purchased from Santa Cruz. Horse-radish peroxidase (HRP)-conjugated goat anti-rabbit IgG polyclonal antibody (cat. no. 1706515) and HRP-conjugated goat anti-mouse IgG polyclonal antibody (cat. no. 1706516) were purchased from Bio-Rad.

### Alkaline comet assay

In 6-well plates, HaCaT cells were seeded at a density of 2 × 10^5^ cells/well and 2 mL of cell culture medium per well. To cause DNA damage, 10 µM Pt was used, and it served as a positive control for DNA damage. The cells were collected and suspended in 100 µL PBS following different treatments. As previously mentioned, the alkaline comet assay was carried out [4]. First, 75 µL of 0.65% regular-melting-point agarose was applied to completely frosted microscope slides, and a coverslip was quickly placed on top. To enable the agarose to harden, the slides were incubated for 3 min at room temperature. After the coverslips were removed, the cell suspension was added to the agarose (1 × 10^6^ cells in 10 µL PBS were combined with 75 µL of 0.65% low-melting agarose). To firm the second coating of agarose, the slides were covered with coverslips once more and put on ice for 3 min. The slides were then submerged in the lysis solution (1 M NaCl, 40 mM EDTA, 4 mM Tris, 14 mM sodium lauroyl-sarcosine, with 1% Triton X-100, pH = 10) for 1 hour at 4 ℃ after the coverslips were removed. After that, the slides were electrophoresed for 20 min at 20 V at 3 A in an alkaline solution (205 mM NaOH, 8 mM EDTA, 5 mM 8-hydroxyquinoline, pH = 13). PBS was used twice to rinse the slides for 5 min at room temperature. A fluorescence microscope was used to view the slides after they had been drained and stained with 200 µL of GelRed at a dilution of 1:10,000. A fluorescent microscope (ECLIPSE Ts2R, Nikon, Japan) was used to capture and analyze cell images. ImageJ version 13.1 software was used to measure the tail moment, olive tail moments, and tail DNA (%).

### Migration assay

In the scratch wound assay, 4×10^5^ HaCaT cells per well were plated in a 12-well plate and incubated until confluence was achieved. The monolayer was then subjected to a scratch using a pipette tip and subsequently washed with serum-free medium to eliminate any detached cells. Following this, the cells were cultured in exosome-depleted complete medium, with or without the addition of Exo^Ctrl^ or Exo^EF^ derived from an equal number of MSCs. Photographs of the cells were taken at 0, 24, 48, and 72 hours post-scratch. The percentage closure of the wound area was calculated using the formula: migration area (%) = (A_0_ - A_n_) / A_0_ × 100, where A_0_ represents the initial wound area and An denotes the remaining wound area at the designated time point.

### Drug treatment

HaCaT cells were planted at a density of 4×10^5^ cells/dish in the galvanotaxis chamber of 100-mm plates to reach approximately 80% density after overnight incubation. Cells were washed twice with PBS. For the MβCD-treated group, cells were treated with fresh serum-free medium containing 4 mM MβCD (cat. no. ST1515-1 g, Beyotime, China) for 1 hour, and then washed twice with PBS and replaced with serum-free medium before the 8 hours DC stimulation started. For the GW4869 group and the LY294002-treated groups, cells were cultured in serum-free medium with 10 µM GW4869 (cat. no. S1971-25 mg, Beyotime, China) or 35 µM LY294002 (cat. no. S1737-5 mg, Beyotime, China) during the 8 hours DC stimulation.

### GPMVs assay

As described previously [21], GPMV buffer (10 mM HEPES, 150 mM NaCl, 2 mM CaCl2, pH = 7.4) was first prepared. DiD (cat. no. C1036, Beyotime, China) and BODIPY-Cholesterol (cat. no. GC42964, GlpBio, USA) were diluted to 2 µM and 1 µM, respectively, and incubated with HaCaT cells for 20 min at 37 ℃ to label cells. Cells were then washed 5 times with PBS and twice with GPMV buffer. Next, for each milliliter of GPMV buffer, 18 µL of 4% Paraformaldehyde and 2 µL of 1 M dithiothreitol solution were added, and cells were incubated in a shaker set to 200 rpm, 37 ℃ for 2 hours to produce GPMVs. The GPMV suspension was transferred to a centrifuge tube and placed at 4 ℃ for 20-30 min to facilitate gravitational sedimentation. Subsequently, the supernatant was carefully aspirated until approximately 200 µL of concentrated suspension remained. The sedimented GPMVs were then resuspended by gentle pipetting and evenly distributed into the galvanotaxis chamber using pipette tips, and the electric field of different strengths and frequencies was applied at 37 ℃ for 2 hours. Finally, the GPMV suspension was collected, and 5 µL of the suspension was added to the center of the microscope slides, and then covered with a 20×20 mm coverslip. Fluorescence was visualized and recorded on a laser scanning confocal microscope (LSM710, Carl Zeiss, GER).

### Statistical analysis

Data are displayed as the mean ± standard error of triplicate unless otherwise noted. A two-tailed Student’s t-test or one-way ANOVA with post hoc tests was used for statistical analysis. Statistical significance was defined as a P value of less than 0.05. GraphPad Prism 9.0 was used to conduct the statistical analysis.

## Acknowledgments

This work is supported in part by Hangzhou Normal University Research Start-up Funds (No. 2019QDL031) and National Natural Science Foundation of China (No. 51807142) to Yan Zhang, and the Scientific Research Fund of Zhejiang Provincial Education Department (No. Y202454801) to Shiwen Huang.

## Author Contributions

Y.Z., S.H., and L.C. designed the experiments, Y.Z., Y.C. supervised the study. S.H. performed the cell experiment and GPMV experiment, S.H. and C.S. did the NTA, L.C. did the TEM, S.H. and L.C. did the analysis. S.H., J. W. and Y.Z. drew the figures, Y.Z., S.H. and L.C. wrote the manuscript. All authors commented and/or edited the manuscript and figures. All authors have given approval to the final version of the manuscript.

---

**Supplementary Figure 1. EFs enhance the production of extracellular vesicles, related to Figure 1**.

(A) Shown are representative phase contrast images of HaCaT cells treated with different strengths of EFs; the direction of EFs is indicated by the black arrow, with the anode on the left and the cathode on the right. (B) The cell viability of each group after 8-hour stimulation was determined by the trypan blue cell counting assay. (C) The average diameter of exosomes produced by HaCaT cells under different EF strengths over 8 hours. (D) Fold changes in the number of MVs produced per cell under different EF strengths over 8 hours. (E) MV number per cell under different conditions. (F) MV number per cell per hour during EF stimulation and post-EF stimulation. (G-J) The diameter distribution of exosomes produced by MSCs and HT1080 cells under different EF strengths over 8 hours. Data are derived from three independent experiments and are presented as means ± SEM. Unless otherwise specified, comparisons were made with the control group. ns indicates not significant, *p < 0.05, **p < 0.01.

**Supplementary Figure 2. EFs facilitate the budding of microscale vesicles into GPMVs, related to Figure 4**.

(A) Confocal microscopy of DiI (red)/BChol (green)-labeled GPMVs exposed to DC/AC EFs (200 mV/mm, 2h) with or without pretreatment with GW4869 revealed spontaneous generation of microscale vesicles. Dynamic membrane fluctuations impede perfect image co-localization. (B) Quantification of microscale vesicle production inside the GPMVs under different conditions. n=30, from three independent experiments. (C, D) The average number of MVs per cell was measured under varying EF stimulations of 200 mV/mm across frequencies ranging from 0 to 400 Hz over 8 hours. AC groups were compared with the DC group. Data are derived from three independent experiments and are presented as means ± SEM. Unless otherwise specified, comparisons were made with the DC EF group. ns indicates not significant, **p < 0.01.

