## Supplementary figures and images for "Electric fields trigger ceramide-dependent budding of exosome vesicles into multivesicular endosomes and boost the generation of exosomes"

### Supplemental Figure 1

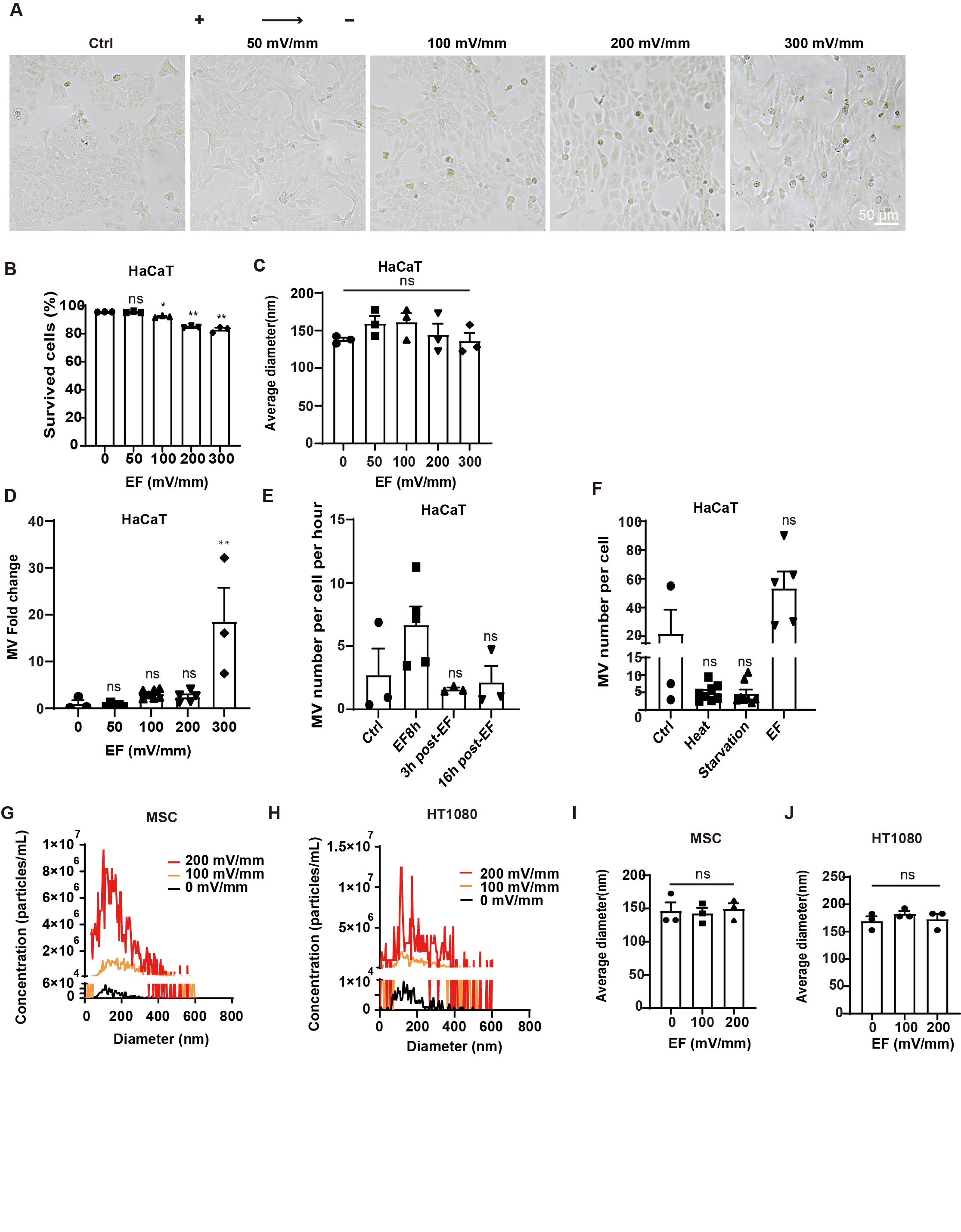

### Supplemental Figure 2

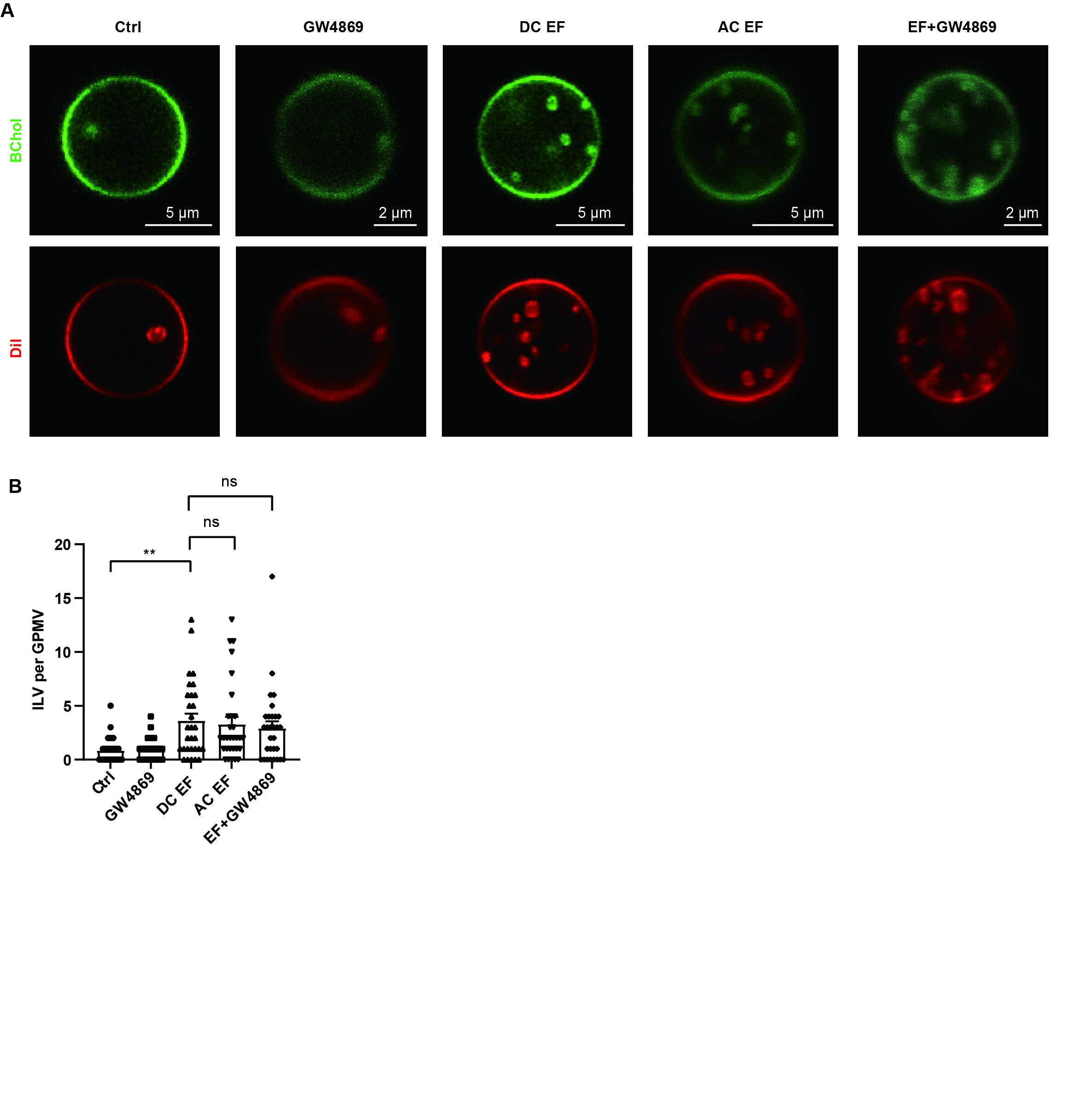
